# Modeling alternative translation initiation sites in plants reveals evolutionarily conserved *cis*-regulatory codes in eukaryotes

**DOI:** 10.1101/2023.10.20.563379

**Authors:** Ting-Ying Wu, Ya-Ru Li, Kai-Jyun Chang, Daisuke Urano, Ming-Jung Liu

**Affiliations:** Institute of Plant and Microbial Biology, Academia Sinica, 11529, Taiwan; Institute of Tropical Plant Sciences, National Cheng Kung University, Tainan, 701, Taiwan; Graduate Program in Translational Agricultural Sciences, National Cheng Kung University and Academia Sinica, Taiwan; Biotechnology Center in Southern Taiwan, Academia Sinica, Tainan 711, Taiwan; Temasek Life Sciences Laboratory, Singapore; Department of Biological Sciences, National University of Singapore, Singapore; Agricultural Biotechnology Research Center, Academia Sinica, Taipei 115, Taiwan

## Abstract

mRNA translation relies on identifying translation initiation sites (TISs) in mRNAs. Alternative TISs are prevalent across plant transcriptomes, but the mechanisms for their recognition are unclear. Using ribosome profiling and machine learning, we developed models for predicting alternative TISs in *Arabidopsis thaliana* and tomato (*Solanum lycopersicum*). Distinct feature sets were predictive of AUG and non-AUG TISs in 5′ untranslated regions and coding sequences, including a novel CU-rich sequence that promoted plant TIS activity, a translational enhancer found across dicots and monocots and also in humans and viruses. Our results elucidate the mechanistic and evolutionary basis of TIS recognition, whereby *cis*-regulatory RNA signatures affect start site selection. The TIS prediction model provides global estimates of TISs to discover neglected protein-coding genes across plant genomes. The prevalence of *cis*-regulatory signatures across eukaryotes and viruses suggests their broad, critical roles in reprogramming the translational landscape in the plant–virus arms race.

**Teaser:** New insights into how plant ribosomes distinguish AUG and non-AUG triplets for protein synthesis via a conserved eukaryotic *cis*-regulatory strategy.

## Introduction

Translation initiation is the first stage of protein synthesis and is also rate limiting, as it includes the selection of the translation initiation site (TIS) in the mRNA. The choice of TIS determines the coding sequence (CDS) of mRNA and ensures the accurate and timely production of a desired protein. This mechanism enables plants to rapidly respond to developmental cues and environmental stress^1,2^. Advanced high-throughput computational and experimental workflows can be used to annotate protein-coding genes, decode plant genomes, and identify the genetic basis of phenotypic diversity among plant species. However, the current criteria for identifying protein-coding genes, which include the presence of an AUG initiation codon, a minimum open reading frame (ORF) length of 100 amino acids, and a (most likely) single ORF in eukaryotic mRNA limit the identification of genes with small or non-AUG-initiated ORFs and may not fully capture the complexity of plant genomes^3,4,5^. Ribosome profiling, which allows the global mapping of TISs *in vivo*, has revealed numerous unannotated TISs in mRNAs in plants^6–10^. These alternative TISs (i.e., the AUG and non-AUG TISs that differ from the annotated AUG sites) mainly located at AUG and near-cognate codons (i.e., codons with one base difference from AUG) direct the translation of uncharacterized ORFs encoding novel peptides/proteins or different protein isoforms. These peptides/proteins play crucial roles in stress and other physiological responses in plants^6–8,10–14^. For example, multiple Arabidopsis (*Arabidopsis thaliana*) small ORFs initiated at AUG encode hormone-like peptides that regulate morphogenic development and salinity stress tolerance^12,15^. The tomato (*Solanum lycopersicum*) valyl-tRNA synthetase gene encodes both mitochondrial and cytosolic proteins via an upstream ACG and annotated AUG initiation site, respectively^6^. Whereas previous annotations of protein-coding genes overlooked alternative TIS-initiated ORFs, ribosome profiling studies have revealed unexpected proteome diversity in plants^6–8,10–14^. Thus, it is crucial to elucidate the general principles of plant TIS recognition to decode plant genome sequences^16^.

How do plant ribosomes recognize start sites for protein synthesis? Although thousands of unannotated AUGs and near-cognate TISs are present in the 5′ untranslated regions (UTRs) and major CDSs of plant mRNAs, ribosomes do not initiate protein synthesis at every triplet they encounter^6–8,10–12^, highlighting the need to understand how start codons and a subset of triplets are selected in mRNA. These mechanisms depend on the sequence context and *cis*-regulatory RNA elements surrounding the start codon^4,17,18^. For example, Kozak sequences and specific TIS-flanking nucleotides in the –2, –4, and +5 positions (where +1 refers to A of the AUG start site) can enhance the efficiency of translation initiation^19,20^. Nevertheless, they do not account for all AUG- and non-AUG TIS activities. Kozak motifs are commonly found in 5′ UTR-nonAUG TISs (TISs located in 5′ UTRs and using a non-AUG codon) and CDS-AUGs TISs (located in CDSs and using an AUG codon) but not in 5′ UTR-AUG TISs (located in 5′ UTRs and using an AUG codon) in both mammals and plants^6,21,22^. Thus, how plant ribosomes recognize different types of start codons in mRNA is not fully understood. Several questions remain, including which sequence features determine alternative TISs in plants, how these determinants jointly and distinctly regulate AUG and non-AUG TIS selection, the similarity of these features across plant species and eukaryotes, and how much of plant genomes truly encodes proteins.

Machine learning (ML) offers one way to uncover the protein-coding genes of plant genomes and understand the mechanisms underlying plant AUG and non-AUG TIS recognition. ML can systematically identify RNA *cis*-regulatory codes of plant alternative TISs and provide more accurate TIS annotations. ML frameworks, which build mathematical models and identify patterns in large datasets, have successfully elucidated complex gene regulatory networks and predicted phenotypes in plants. For example, ML-based predictions of dynamic gene expression responses and identification of novel regulators have revealed the transcriptional regulatory network architectures of plant stress responses^23–25^. ML strategies have also helped characterize novel *cis*-regulatory DNA elements and their combined effects in regulating transcription and associating genetic variation with differential gene expression and phenotypic diversity^26,27,28,29^. Therefore, integrating high-throughput translation initiation sequencing datasets from different plant species using ML techniques may facilitate TIS/ORF annotations and help decipher the underlying TIS-determining principles across plants^6,10^.

Here, we combined ML-based, computational, and experimental techniques to investigate how plant ribosomes recognize different types of TISs along mRNA sequences, including those initiated at both canonical AUG and non-AUG codons, and explored uncharacterized TIS-initiated ORFs in Arabidopsis and tomato. We built ML models using TIS-flanking mRNA sequence and TIS codon usage and characterized common or species-specific sequence features including the novel CU-rich sequence features. We explored their conserved regulatory role across dicots and monocot plants, viruses, and humans. Our findings provide a global view of alternative TISs in genomes, paving the way for improved gene annotation in plants. The novel CU-rich features we found not only shed light on the regulatory mechanisms of alternative TIS usage across plants and in eukaryotes but also deepen our understanding of virus protein expression strategies in hosts.

## Results

### Machine learning enables cross-species TIS predictions in Arabidopsis and tomato

To comprehensively and precisely build models that predict TISs in plant mRNAs, we utilized Arabidopsis and tomato ribosome profiling datasets^6,10^ to globally profile experimentally supported alternative AUG and near-cognate TISs (referred to as true-positive [TP] TISs) and implemented an ML workflow to distinguish these TP TISs from AUG and near-cognate triplets with no significant translation initiation signals (true-negative [TN] TISs) (Fig. 1a–d). Bioinformatics and statistical analyses identified TPs (see Methods) and categorized them into six groups based on the locations of initiation codons and their sequences: 5′ UTR-AUG, 5′ UTR-nonAUG, CDS-AUG, CDS-nonAUG, 3′ UTR-AUG, and 3′ UTR-nonAUG (left panel, Fig. 1a). We identified several hundred to thousands of TP TISs in the 5’ UTR and CDS but not the 3′ UTR, which is mainly located at AUG and near-cognate codons in both Arabidopsis and tomato (Fig. 1b, c; Extended Data Fig. 1). Therefore, we generated TN TIS sets of the triplets at the AUG and near-cognate codons with no significant ribosome profiling signals and upstream of the TP TISs (see Methods; Fig. 1a-c) and focused on the TP and TN TISs in the 5′ UTR and CDS for further analysis using the ML workflow illustrated in Fig. 1d. First, we collected 2657 features, comprising 8 known (Kozak and previously reported nearby flanking sequences^6,19,20,30–32^), 23 ORF (mononucleotide contents and secondary RNA structures upstream of or within ORFs and ORF sizes), and 2626 contextual features (nucleotide/amino acid frequency of k-mers in the 200-nt region centered on a TIS) for each TIS (right panel, Fig. 1a) and generated a balanced dataset with equal numbers of TPs and TNs by random sampling (Fig. 1d, step 1, see Methods). Second, we imputed and scaled the feature data to make them comparable. Third, we selected the final feature set for training (70%) and testing (30%) data based on the correlation between features and the significance of enrichment between TPs and TNs, as too many features can interfere with prediction performance^33^ (Fig. 1d, step 2, see Methods). Fourth, we generated ML models using four algorithms and compared their performance to assess the predictive power of different feature categories (Fig. 1d, step 3, see Methods).

**Figure 1.**
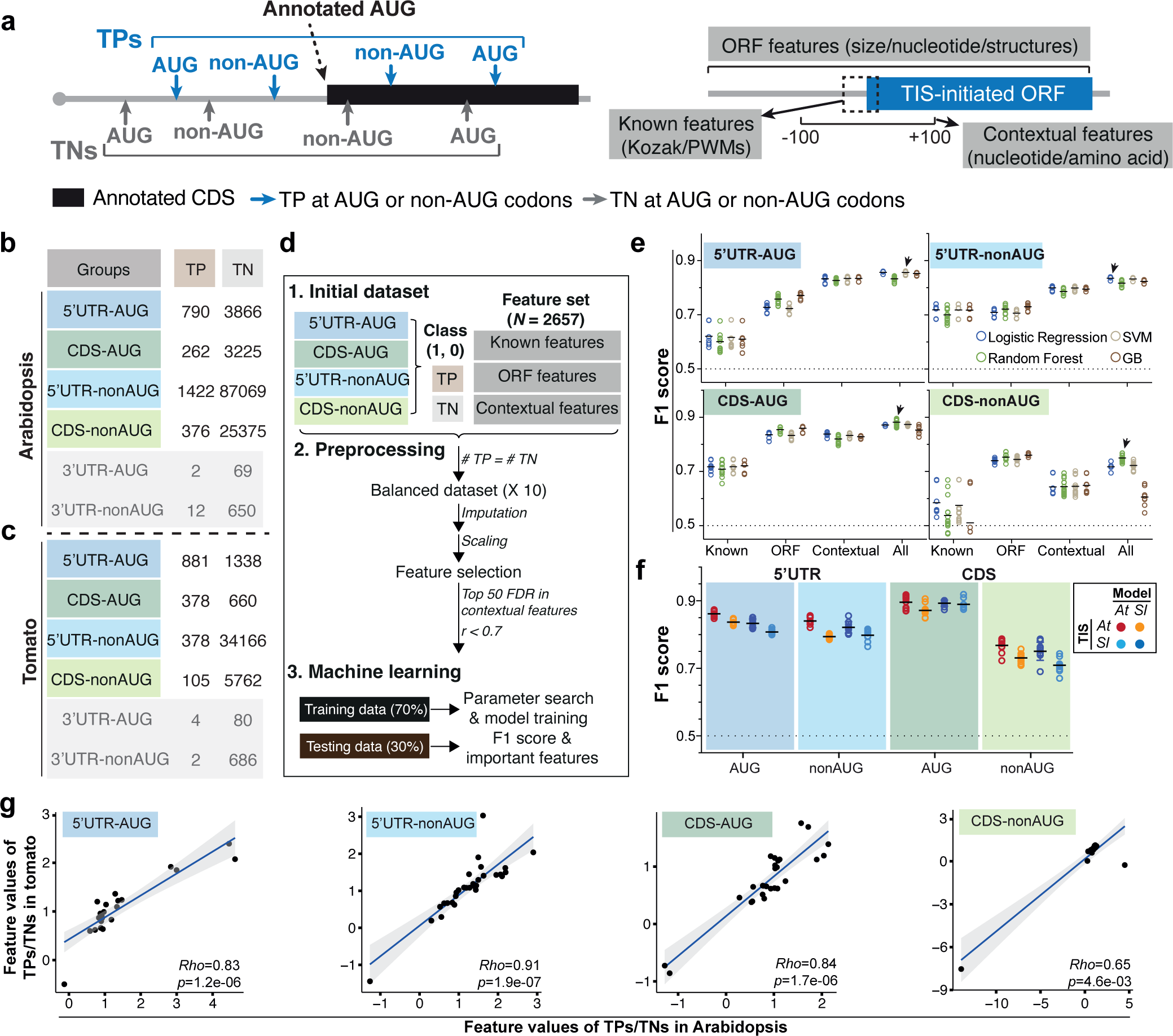
Identification of prediction models and the associated features that predict Arabidopsis and tomato alternative translation initiation sites (TISs). (a) Illustration of the identified true positive (TP, blue) and true negative (TN, gray) TISs that were categorized based on the location of initiation codons (i.e., 5′ UTR, CDS, or 3′ UTR) and their sequences in a transcript. The “Known” features include the Kozak motif and position weight matrixes (PWMs) generated from short sequences centered on TISs^30,67^. The “ORF” features consist of various nucleotide compositions/sizes of the alternative TIS-initiated open reading frames (ORFs) and the annotated ORFs and also the RNA structures/nucleotide compositions of their surrounding regions. The “Contextual” features based on *K*-mer enrichment analyses (*K*=1– 3) consist of the nucleotide and amino acid sequence contexts in the 200-bp region centered on the TISs (See Methods). Gray line, black and blue boxes: mRNAs, annotated AUG TIS- and alternative TIS-initiated ORFs. (b,c) Numbers of the identified TP and TN TISs, categorized as described in (a), in Arabidopsis (b) and tomato (c). (d) Machine-learning (ML) workflow used to identify prediction models and the features that were informative for predicting TISs (See Methods). Pearson correlation coefficient *(r)* and the false discovery rate (FDR; determined by Wilcoxon-rank sum test) represent the correlation and the statistical significance of differences for the feature values between TP and TN TIS sets. “Top 50 FDR in contextual features” represented that the 50 contextual features with smallest FDRs were selected for further analysis (See Methods). F1 score: the harmonic mean of precision and recall, which ranges from 0 to 1 with 1 indicating a perfect model. (e) The F1 scores showing the performances of four different ML algorithms based on different sets of features in predicting the four Arabidopsis TIS groups (i.e., the AUG and non-AUG TISs located in the 5′ UTR and CDS). Circle: the performance of a model on a randomly balanced TP and TN TIS dataset; black line: mean of the F1 scores for a given ML algorithm. Arrow: the best model (i.e., for the ML algorithm with highest mean performance, the model with highest F1 score); dashed line: the baseline performance expected by random guessing. (f) The cross-species and within-species prediction performance (shown as F1 score) when using the best model built in one species to predict the TISs in another species (light blue and orange) and to predict the TISs within the same species (red and dark blue). Results are shown for the four TIS groups in Arabidopsis and tomato. (g) Rho: Spearman’s rank correlation coefficient. The black line indicates the fitted linear regression line, and the gray area indicates the 95% confidence interval.

Focusing on Arabidopsis TISs, the model employing the features from all categories outperformed those employing known, ORF, or contextual categories for TIS prediction (Fig. 1e). The F1 scores (representing prediction performance) ranged from 0.7 to 0.9, with the highest and lowest F1 scores observed for the 5′ UTR-AUG and CDS-nonAUG groups, respectively (arrows, Fig. 1e; Supplementary Table 1). Therefore, in addition to using features with known biological functions (e.g., Kozak score), using a combination of unexplored features (e.g., ORF and contextual features) and features with known biological functions is important for TIS prediction.

To assess the generality of TIS recognition mechanisms across plants, we explored whether the models generated from Arabidopsis could be used for predictions in tomato. We generated the best models that predicted the four types of tomato TISs with F1 scores ranging from 0.7 to 0.88 (dark blue, Fig. 1f; Extended Data Fig. 1e and Supplementary Table 1), showing the robustness of the established ML workflow in identifying TIS prediction models across plants. Whereas the Arabidopsis model performed slightly worse when used to predict the tomato TIS dataset for all four groups (light blue vs. dark blue; Fig. 1f), cross-species prediction performed well, with F1 scores > 0.7 for most groups (Fig. 1f). We observed similar patterns using tomato models to predict Arabidopsis TISs (red vs. orange; Fig. 1f). The feature enrichment values (ratio of TP to TN feature values) showed significant positive correlations in Arabidopsis and tomato for all groups tested (Fig. 1g). These results highlight the conservation of TIS prediction models and the similarity of significant features across dicot plants, suggesting that information gained from one species-based model could be useful for predicting TISs in other plant species.

### Distinct nearby sequence contexts and codon usage bias are associated with plant AUG and non-AUG TIS predictions

We asked to what extent each feature contributed to the performance of the TIS prediction model in Arabidopsis and tomato (Fig. 2). Due to the relatively poor performance (F1 score = 0.73–0.75) of CDS-nonAUG prediction in both species, we focused on the three other groups (F1 scores = 0.81–0.88; Fig. 1e; Extended Data Fig. 1e and Supplementary Table 1). We identified the top features that contributed most to model performance for each Arabidopsis TIS group (Extended Data Fig. 2a). We used the known feature of PWM-5′ UTR-TP (position weight matrix [PWM]), representing the nucleotide compositions of the flanking regions (positions –15 to +10) for 5′ UTR-AUG and 5′ UTR-nonAUG TP TISs, to calculate the sequence similarity of the TIS-flanking regions between the target TIS and the 5′ UTR TPs, as reported previously ^6,20,30,31^. The PWM-5′ UTR-TP feature substantially contributed to the accuracy of the model, especially for predicting Arabidopsis 5′ UTR-nonAUG TISs (Fig. 2c,d), as supported by its high feature importance score and significant enrichment (false discovery rate [FDR], as determined by Wilcoxon signed-rank test with Bonferroni correction; Fig. 2c), its top feature ranking and higher feature occurrence in 10 randomly balanced TP/TN datasets (right panels; Fig. 2d), and the differential feature values between TPs and TNs (left panels; Fig. 2d).

**Figure 2.**
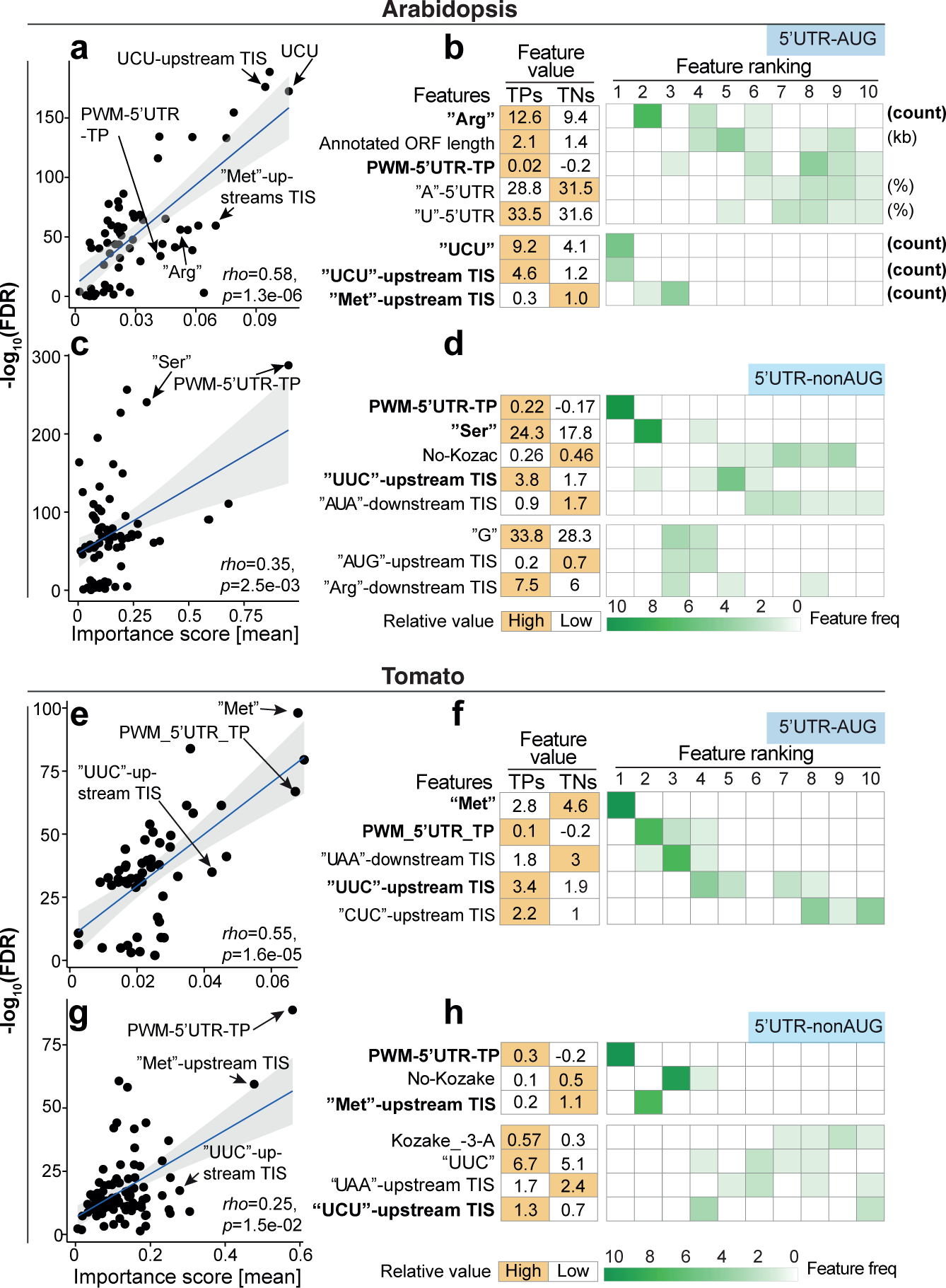
The features that were most informative for predicting 5′ UTR TISs in Arabidopsis and tomato. (a) Comparison of the importance scores derived from the model and the statistical significance of differences (-log_10_(FDR), determined by Wilcoxon signed-rank test) between Arabidopsis 5′ UTR -AUG TPs and TNs for the features used in the best model. Rho: Spearman’s rank correlation coefficient. The black line indicates the fitted linear regression line, and the gray area indicates the 95% confidence level interval. (b) The means of the feature values in the Arabidopsis 5′ UTR - AUG TP and TN datasets (right) and the frequency of features identified in 10 randomly balanced datasets (left) for the Feature Elimination-determined top10 features (ranked using their importance; see Extended Data Fig. 2a). The rank and frequency indicate the importance of a given feature in the prediction model and their robustness using 10 randomly balanced datasets. The features with frequency >7 within the top 10 are shown. Orange indicates the TIS group with the higher feature value. (c-h) As described in (a,b), but for the Arabidopsis 5′ UTR -nonAUG TIS group (c,d), the tomato 5′ UTR -AUG TIS group (e,f) and tomato 5′ UTR -nonAUG TIS group (g,h).

Analysis of the nucleotide compositions of the TIS-flanking regions showed that all three Arabidopsis TIS groups have a high frequency of C nucleotides in nearby flanking regions (Fig. 3a; Extended Data Fig. 3a). By contrast, only the 5′ UTR-nonAUG and CDS-AUG TPs tended to have As at positions –4 to –1 and G and C at positions +4 and +5, respectively (Fig. 3a; Extended Data Fig. 3a). This sequence-specific pattern of 5′ UTR-nonAUG and CDS-AUG TPs, but not 5′ UTR-AUG TPs, was similar to that of annotated AUG TPs (Fig. 3a,c; Extended Data 3a,c) and is in line with previously reported sequence requirements for mammalian TISs at AUG and non-AUG codons^20,31^. We observed similar patterns when examining the prediction models of tomato TISs (Figs. 2e–h, 3b,d; Extended Data Figs. 2b,g–j and 3b,d), reflecting the similarity of sequence features between Arabidopsis and tomato (Fig. 1g). These observations suggest that 5′ UTR-AUG, 5′ UTR-nonAUG, and CDS-AUG TISs have both shared and distinct sequence context dependencies for TIS recognition in plants.

**Figure 3.**
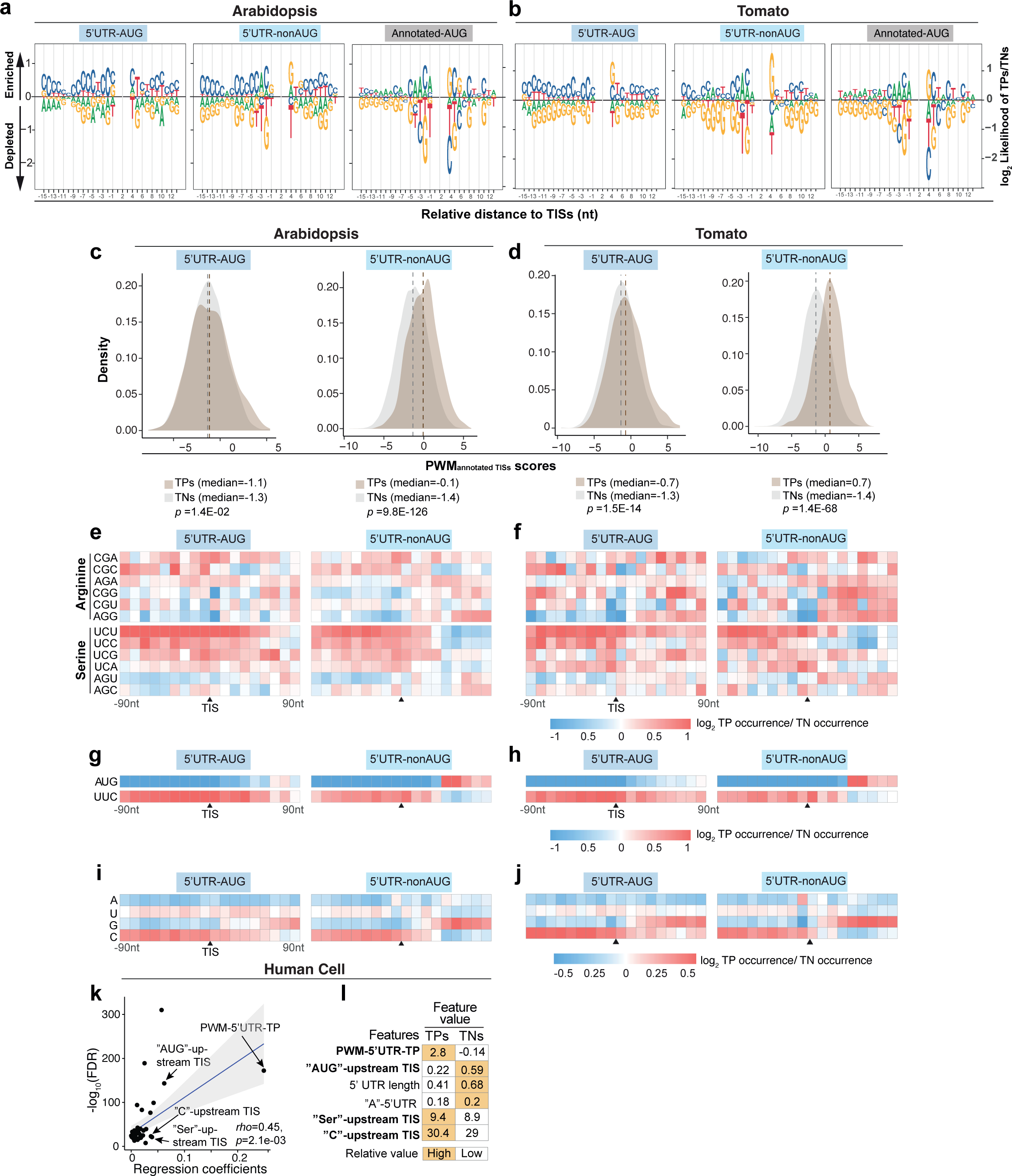
The C/U nucleotide compositions and the flanking sequences of Arabidopsis and tomato 5′ UTR TISs. (a,b) Sequence logo plots showing the differential enrichment of A/U/C/G nucleotides between TPs and TNs in the regions 15-bp upstream and 13-bp downstream of the TISs, represented as the log2 ratio of the site frequencies between TPs and TNs, for the 5′ UTR TISs and annotated AUG sites in Arabidopsis (a) and tomato (b). (c,d) Position-weight matrix (PWM) scores showing the sequence similarity of the TIS-flanking regions between TPs (brown)/TNs (gray) and annotated TISs for the Arabidopsis (c) and tomato 5′ UTR TISs (d). PWM_annotated TISs_ matrix was determined based on the annotated TISs with *in vivo* translation initiation activities (See Methods). *P*-values derived from Mann–Whitney U test were used to evaluate the significance of differences in PWM scores between TP and TN datasets. Dashed line indicates the median value. (e,f) Enrichment of sites with the indicated 3-mer sequences in Arabidopsis (e) and tomato (f) TIS groups, represented as the log2 ratio of the site frequencies between TPs and TNs in the 180-bp region centered on Arabidopsis TISs with a 10-bp window. (g-j) As described in (e,f), but for the AUG and UUC sequences (g,h) and the A, U, C, and G mononucleotides (i,j). (k) As described in Fig. 2a, but shown for the regression coefficients (x-axis) for the features used in the best linear regression model of predicting human TISs. (l) As described in Fig. 2b, but for the top 6 features with highest regression coefficients.

The model performances were slightly worse for the non-AUG relative to AUG groups regardless of TIS location in both Arabidopsis and tomato (Fig. 1e; Extended Data Fig. 1e and Supplementary Table 1). Not every near-cognate codon serves as a TIS with equal activity, with CUG, ACG, and GUG generally being the most efficient^4,6,31^. Hence, we attempted to increase prediction accuracy by adding the feature TIS codon usage bias (i.e., enrichment values of each near-cognate codon among TP TISs). In general, models with TIS codon usage bias information slightly outperformed those without this feature (Extended Data Fig. 4a), and the rate of TNs→ TPs misclassification (i.e., a TN TIS inaccurately classified as a TP TIS) decreased by 6%–10% (light brown, Extended Data Fig. 4b). The major codons contributing to misclassification were AAG and AGG in both species (arrows, Extended Data Fig. 4c), suggesting that, in addition to the flanking sequences context, the codon preference of TISs themselves is also important for non-AUG prediction and that this feature is evolutionarily conserved across plants.

### CU-rich tracts are important for plant TIS predictions

In addition to known sequence features, we also identified several ORF/contextual features that are important for model accuracy, such as “UCU” short sequences, with the highest importance score for predicting Arabidopsis 5′ UTR-AUG TISs (Fig. 2a). This result is consistent with the observation that the model employing features from all categories outperformed the model employing known features (Fig. 1e). We explored the ORF/contextual features that strongly contributed to model accuracy in each Arabidopsis and tomato TIS group: the codons encoding serine (Ser) and methionine (Met), the short sequences “UUC” and “UCU,” and the mononucleotides (e.g., A, U, C, or G) (indicated by arrows in Fig. 2a,c,e,g; bold in Fig. 2b,d,f,h). Since “UCU” is one of six codons encoding serine, we examined the degrees of enrichment of all six Ser-encoding codons in the TIS-flanking region between TPs and TNs. “UCU” and “UCC” were more enriched than the other codons (“UCU”, “UCC,” and “AGC” medians = 0.62∼1.1, 0.41∼0.7, and –0.06∼0.07 in Arabidopsis 5′ UTR-AUG, 5′ UTR-nonAUG, and CDS-AUG TIS groups, respectively; Fig. 3e; Extended Data 4a). In addition, “UUC” was present at higher frequency and “AUG” was present at lower frequency in the sequences surrounding TP TISs than TN TISs (Fig. 3g; Extended Data 4c). Analysis of individual mononucleotides (A, U, C, or G) showed that “C” is highly enriched in the peripheral sequences near TISs (Fig. 3i). We observed similar patterns for tomato TISs (Fig. 3f,h,j; Extended Data Fig. 5b,d,ef). Intriguingly, in the best model for predicting human TISs generated in a previous study^30^, the features “C content” and “Ser frequency” strongly contributed to prediction power (indicated by arrows, Fig. 3k) and showed significantly differential feature values between TPs and TNs in the upstream regions of 5′ UTR-AUG and 5′ UTR-nonAUG TPs (bold, Fig. 3k,l). These results suggest that CU-rich nucleotide tracts, regardless of the amino acids they encode, are important for alternative TIS prediction and are a shared *cis*-regulatory RNA signature across eukaryotes.

Contextual features included the 1-to 3-nt sequences surrounding TISs (see Methods). We asked whether longer specific sequences, representing putative *cis*-regulatory RNA elements^1,34^, also regulate TIS selection during ribosome scanning of mRNAs. We used a previously established enrichment-based *k*-mer (k>3)-finding pipeline to identify significantly enriched sequences^26^. The identified *k*-mers (most ranging from 4–6 nt) alone accurately predicted Arabidopsis TISs for all groups (Fig. 4a; Supplementary Table 1), especially for the 5′ UTR-AUG and CDS-AUG groups, and showed prediction performances comparable to that of all (known/ORF/contextual) features (Fig. 1e). The “UCUUC” and “UCUCU” sequences played substantial roles in model prediction, as supported by their highest importance scores (orange, Fig. 4b; Extended Data Fig. 5g) and more frequent occurrence in TPs (blue, x-axis in Fig. 4c; Extended Data Fig. 5i), respectively, and their predominant enrichment in the upstream regions of TP TISs (Fig. 4e; Extended Data Fig. 5i). There was also a positive correlation between the number of C/Us and the enrichment of sequence occurrence of all 5-mers for all TIS groups (Fig. 4c; Extended Data Fig. 5i), supporting the observation of C/U mono/trinucleotide enrichment (Fig. 3e–j). The tomato TP TISs showed similar tendencies (Fig. 4d,f; Extended Data Fig. 5h,j). These results suggest that both CU-rich nucleotide tracts and specific CU-related putative *cis*-elements are associated with alternative TIS selection in plants.

**Figure 4.**
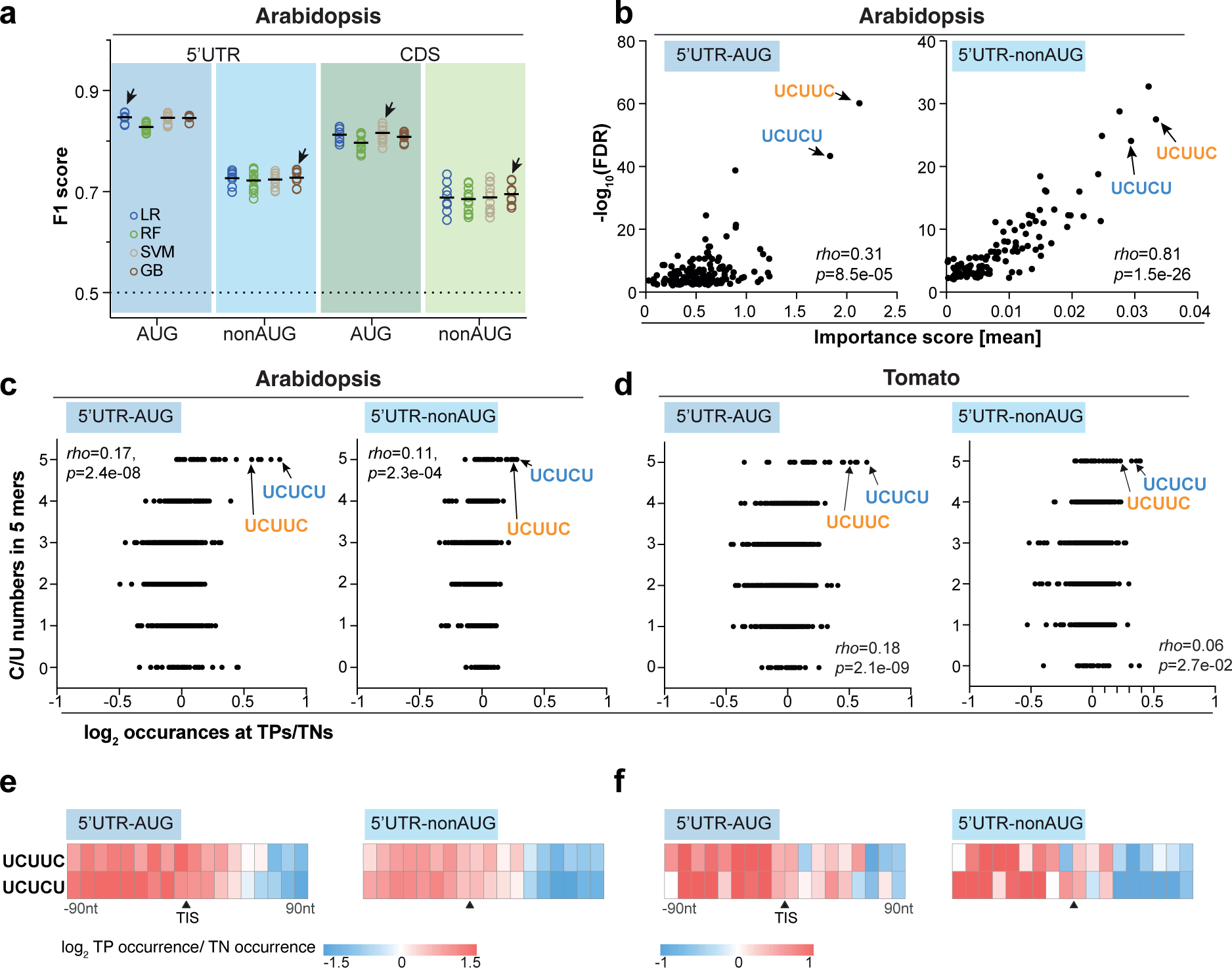
Characterization of putative RNA cis-elements predictive of TISs in Arabidopsis. (a) As described in Fig. 1e, but for the model employing the identified putative RNA *cis*-elements to predict the four Arabidopsis TIS groups. Arrows: the best model. (b) As described in Fig. 2a, but for the putative RNA *cis*-elements in predicting the Arabidopsis 5′ UTR -AUG and 5′ UTR - nonAUG TIS groups. (c,d) Relationship between the number of C/Us (y-axis) and the enrichment of sequence occurrence (x-axis) for all 5-mers in the Arabidopsis (c) and tomato (d) 5′ UTR -AUG and 5′ UTR -nonAUGTIS groups. The enrichment is represented as the log_2_ ratio of median sequence occurrence between TPs and TNs in the 200-bp regions centered on TIS sites. Rho: Spearman’s rank correlation coefficient. (e,f) As described in Fig. 3E, but for the enrichment of putative RNA *cis*-elements in Arabidopsis (e) and tomato (f) 5′ UTR TIS groups. The putative “UCUUC” and “UCUCU” elements with the highest importance score and with the highest enrichment, which are indicated by arrows and highlighted in blue and orange in (b,c,d), are shown.

### The plant CU-rich tracts function as translation enhancers to promote initiation activity

To assess the roles of the CU-related features identified by ML in recognizing non-canonical TISs, we mutated the CU-rich nucleotide tracts (by changing C to G when C was flanked by U) in the upstream 100-nt regions of selected alternative AUG and non-AUG TISs reported previously^6^ and identified in this study (see Extended Data Fig. 6a–d for details). In three of the four TISs examined, mutating CU-rich tracts led to much lower protein abundance (WT vs. mCU [CU-tract mutation]; Fig. 5a), whereas steady-state mRNA levels remained comparable (Fig. 5b), indicating that mutating the CU-rich tracts generally decreased the efficiency of protein synthesis. Mutating “UCUUC” and “UCUCU” in the 5′ UTR-AUG TIS of Solyc06g076770.3.1 also decreased protein abundance (WT vs. mUCUUC/mUCUCU [UCUUC/UCUCU mutations]; Fig. 5a). Note that mutating either the CU-tract or “UCUUC”/”UCUCU” sequences in the 5′ UTR-nonAUG TIS of Solyc06g009750.3.1 did not affect translation efficiency (Fig. 5a,b), suggesting that additional parameters contribute to TIS activity.

**Figure 5.**
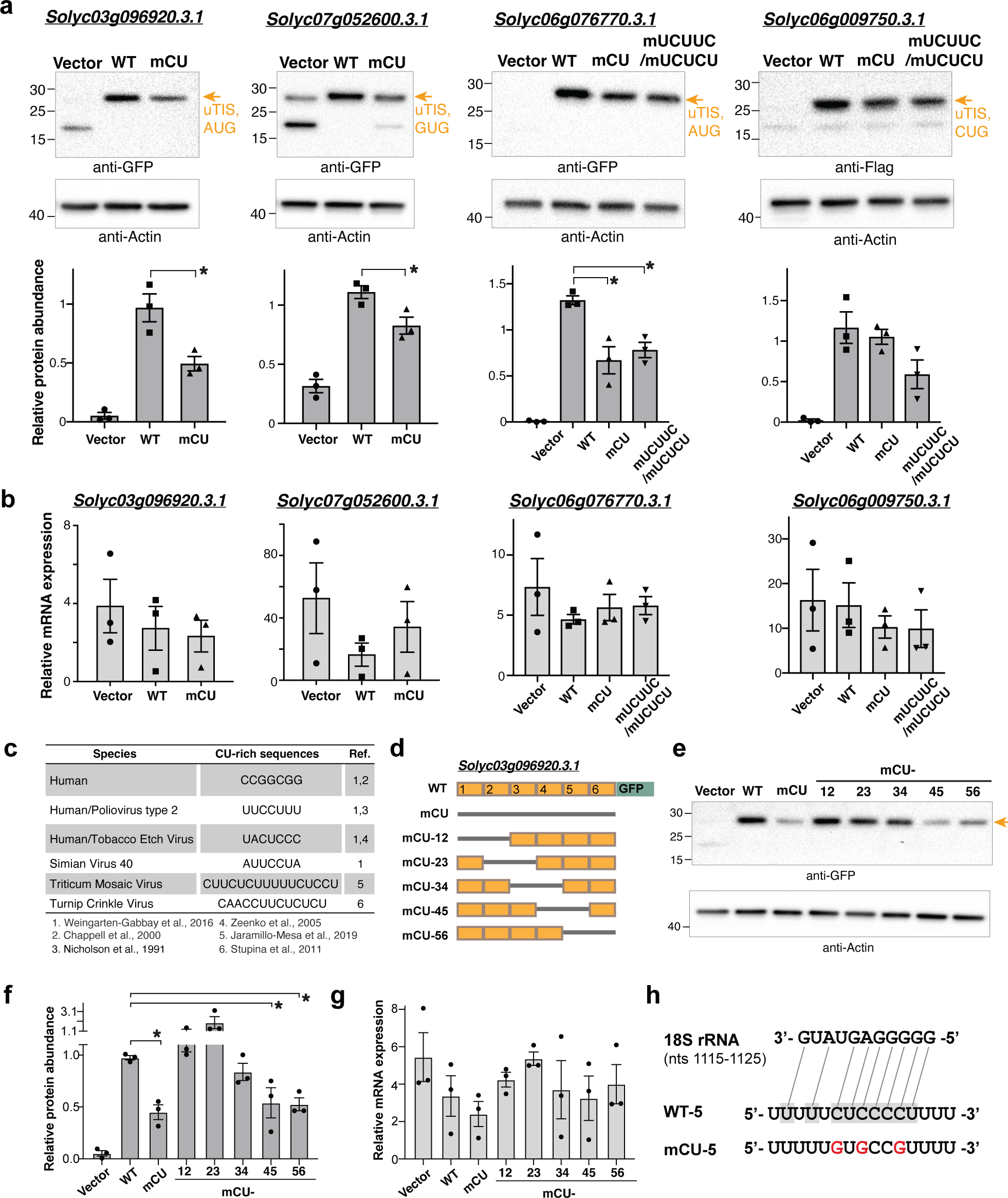
Mutation of CU-rich sequences on plant cellular mRNAs attenuated the TIS activities. (a) Immunoblotting analyses of proteins with translation initiated from the TISs indicated in Extended Data Fig. 6 (orange arrows) and with translation driven by the upstream 100-nt wild-type (WT) sequence or sequences with mutations of CU tracts (mCU) or UCUUC/UCUCU (mUCUUC/mUCUCU) sites. Proteins were expressed in *Nicotiana benthamiana* (tobacco) leaves using the Agrobacterium-mediated transient expression system. Vector: tobacco leaves infiltrated with agrobacteria containing the expression vector (i.e., the GFP or Flag-containing plasmid without a target gene sequence). The protein abundance of the reporter genes relative to Actin for three biological repeats (dots) and the corresponding means and standard errors are shown (bottom panels). (b) As indicated in (a), but shown for the mRNA abundance of the TIS-containing transcripts relative to the *UBQ3* in transformed plants in (a) determined by quantitative RT-PCR analyses. (c) The known CU-rich sequences found on human transcripts and on plant and animal viral mRNAs promote translation efficiency as reported in the indicated literatures. (d) Illustration of swapping the Wild-Type (WT, orange box) and the CU-mutated (mCU, gray line) sequences for the TIS-upstream region of a 5′ UTR -AUG TIS of Solyc03g096920.3.1 examined in (a). The TIS-upstream region was divided into six subregions with the sizes of 15∼17 nts (indicated in Extended Data Fig. 6a) to generate distinct mCU mutants with indicated CU-mutated regions fused with the GFP reporter gene as described in (a). (e,f,g) As described in (a,b), but for the protein (e,f) and mRNA abundances (g) of the mCU mutants with fully or partially mutated regions as indicated in (d). (h) The putative binding site on WT sequences (shaded), but not on CU-mutated ones, to the plant 18S rRNA (sold and dashed lines). The sequences of the plant 18S rRNA at positions nt 1115 to 1125 and the wild-type sequence or sequence with mutations of CT tracts (red) in the 5^th^ region indicated in (d) are shown. *: *p*-values <0.05, which is derived from one-way Anova test, representing the significant difference between WT and the samples with CU sequence mutations.

Specific CU-rich elements enhance TIS activity in an internal ribosome entry site (IRES)–like manner in human, human viral, and plant viral mRNAs (Fig. 5c)^35–40^, whereas their roles in plants are largely unexplored. To characterize the CU-rich elements in plant mRNAs, we mapped the CU-rich sequences that predominantly govern translation initiation efficiency. We swapped the TIS upstream regions between WT and mCU mutants from the 5′ to 3′ end in the 5′ UTR-AUG TIS of Solyc03g096920.3.1 and assessed the effect on translation efficiency of a green fluorescent protein (GFP) reporter (Fig. 5d). GFP protein signals were significantly weaker for mCU-45 and mCU-56 than for the WT and were most similar to that of mCU (Fig. 5E,F); however, their mRNA abundances were comparable (Fig. 5g). The mutated 5^th^ region was present in both mCU-45 and mCU-56, pointing to the role of the CU-rich sequence in this region in regulating translation initiation. Supporting this notion, the WT 5^th^ regions contained a CU-rich site that can base-pair with a purine-rich (especially G) region of plant 18S rRNAs, a highly conserved region across rice (*Oryza sativa*), tobacco (*Nicotiana tabacum*), maize (*Zea mays*), and wheat (*Triticum aestivum*) that efficiently enhances translation^41^. These results provide experimental evidence that CU-rich elements enhance translation initiation in plant mRNAs, likely by increasing complementary interactions between mRNAs and 18S rRNA within 40S ribosomes^41^.

The findings of the ML-characterized importance of CU-rich features in Arabidopsis, tomato, and human mRNAs (Figs. 2,3,4) and the experimentally supported role of specific CU-rich elements in translation initiation in human, viral, and plant mRNAs (Fig. 5) point to a conserved translational strategy of CU-rich tracts and specific CU-related *cis*-elements in regulating AUG and non-AUG TIS recognition across plants and even in eukaryotes and viruses.

### TIS prediction models discover plant TISs

Although our TIS prediction models performed well (Fig. 1; Extended Data Fig. 1e), ∼9%– 25% of TNs were misclassified as TPs (i.e., TNs with a high TP prediction score and classified as TPs; TNs→TPs) in each group (light brown in Fig. 6a,b; Extended Data Fig. 7a,b). Note that the Arabidopsis and tomato TN sites were identified based on sequencing signals from ribosome profiling, which were generated from Arabidopsis suspension cells and tomato leaves under a single condition, respectively (Fig. 1)^6,10^. Experimental and/or analytic limitations, including inadequate sequencing depth, low sample diversity across plant tissues/stages, and the presence of specific translation inhibitors^42–44^ might reduce the accuracy of identifying TN triplets and thus lead to misclassified TNs (TNs→TPs). Thus, to explore the possibility that misclassified TNs function as initiation sites, we compared the feature values between correctly predicted TNs (true TNs; TNs→TNs) and incorrectly predicted TNs (misclassified TNs; TNs→TPs) in Arabidopsis (Fig. 6a; Extended Data Fig. 7a). The features that likely contributed to misclassification (i.e., with significant enrichment [FDR] and high frequency in 10 randomly balanced TP/TN datasets; indicated by arrows in Fig. 6c; Extended Data Fig. 7c) were those with high importance scores in the built prediction models (Fig. 2; Extended Data Fig. 2)^45^. For example, in the Arabidopsis 5′ UTR-nonAUG group, the number of Ser residues (which reflects CU nucleotide enrichment) was significantly different between mispredicted TNs (TNs→TPs) and true TNs (TNs→TNs) (Fig. 6c), and this feature also had a high importance score in the corresponding model (Fig. 2d, “Ser”). We assessed the values of these features (e.g., number of Ser residues in the 5′ UTR-nonAUG group) among the mispredicted and correctly predicted groups. Indeed, the features important for TN misclassification (TN→TP) all had values similar to those of TPs (TP→TP) (Fig. 6e). These patterns were also observed for misclassified tomato TISs (Fig. 6b,d,f; Extended Data Fig. 7b,d,f).

**Figure 6.**
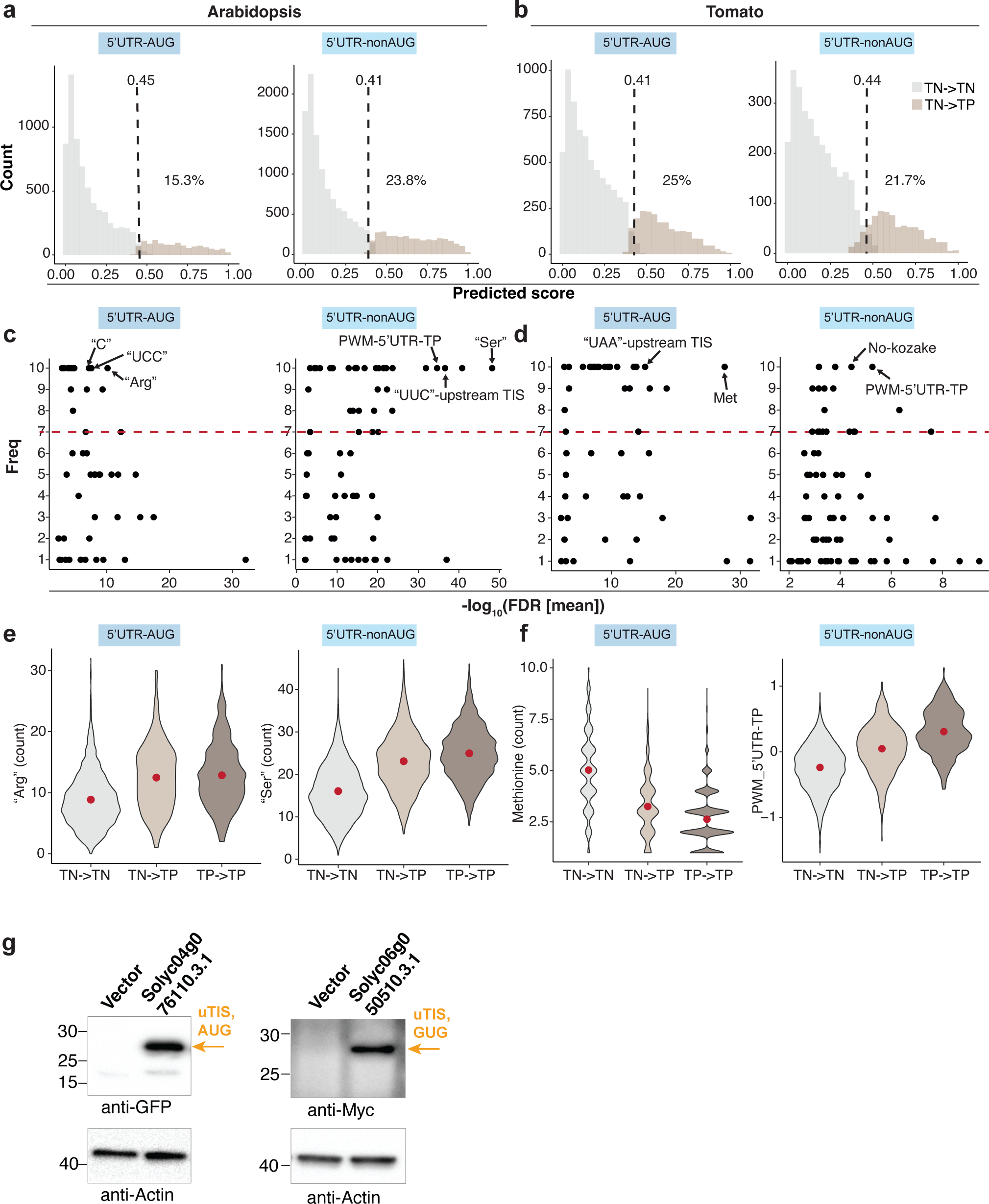
The features of mis-classified TN TISs. (a,b) Prediction score distribution of the TN TISs for the 5′ UTR-AUG and 5′ UTR-nonAUG TIS groups in Arabidopsis (a) and tomato (b). The mean threshold (dashed lines) for the classification of TN→TN (the TN TISs predicted as TNs, gray) and TN→TP (the TN TISs predicted as TPs, light brown) and the mean percentage (%) of the TN TISs mis-classified as TP (TN→TP group) derived from the best models in the10 randomly balanced datasets (as indicated in Fig. 1) are shown. (c,d) Dot plots show the frequency (y-axis) of a given feature used for TIS prediction in 10 randomly balanced datasets and the feature enrichment (FDR, x-axis) between the TN→TN and TN→TP groups for the TIS groups indicated in (a,b). The red line represents the threshold (frequency ≥ 7) of important features as indicated in Fig. 2. (e,f) Violin plots show the feature value distributions for the features of “Arg”, “Ser” and methionine counts or PWM-5′ UTR-TP that were most enriched in (c,d) for the TN→TN (gray) and TN→TP (light brown) groups, indicated in (a), and the TP→TP (the TP TISs predicted as TPs, dark brown) group. The red dot represents the median value. (g) As described in Fig. 5a, but for the Immunoblot analyses of proteins translated from the misclassified TISs shown in Extended Data Fig. 6. Vector: tobacco leaves infiltrated with agrobacteria containing the expression vector (i.e., the plasmid without a target gene sequence).

To explore the potential of the TIS prediction models to identify alternative TISs in plant mRNAs, we assessed whether the mispredicted TNs with high prediction scores function as TISs *in vivo.* These TISs included those with a low read count (the 5′ UTR-AUG TIS in the MYB transcription factor gene Solyc06g076770.3.1 and 5′ UTR-CUG TIS in Solyc06g009750.3.1, encoding an importin alpha involved in nuclear transport); a low precision of ribosome-protected fragments mapping to the TIS (5′ UTR-AUG TIS in Solyc04g076110.3.1, encoding a protein of unknown function); or no read signals (5′ UTR-GUG TIS in Solyc06g050510.3.1, encoding a DNA repair protein) (details in Extended Data Fig. 6c–f). Immunoblotting detected proteins corresponding to the expected sizes for these misclassified AUG- and nonAUG TIS-initiated ORFs (vector vs. WT; Figs. 5a,6g). In addition, the mispredicted CDS-AUG TIS site of Solyc11g039830.2.1 (encoding Glycyl-tRNA synthetase) could potentially generate a protein isoform with distinct N termini, likely affecting mitochondrion targeting signals (Extended Data Fig. 6g). Therefore, alternative TISs can be located in genes with known functions and direct the translation of novel polypeptides (thereby increasing the coding potentials of genes) or protein isoforms that diversify the organelle proteome. Thus, TIS prediction models can help identify potential TISs, even without experimental evidence.

### Predicting TISs with conserved features in monocots and dicots via transfer learning

To explore whether our TIS prediction models can be applied to different tissues or even monocots, we generated TIS lists from the cycloheximide-treated ribosome profiling datasets of different plant species and tissues^6,10,46,47^ using RiboTISH software^48^. We grouped the RiboTISH-reported TISs based on FDR value (i.e., the adjusted *p*-value of frame test provided by RiboTISH) and examined their prediction scores generated by our best Arabidopsis TIS prediction model (Fig. 1e). Compared to TISs with lower FDRs (gray and light green lines; top and middle panels in Fig. 7a; Extended Data Fig. 8), TISs with higher FDRs had higher prediction scores in three types of TIS groups, especially 5′ UTR-nonAUG, in Arabidopsis (suspension cells) and tomato (leaves) (lime green and dark green lines; top and middle panels in Fig. 7a; Extended Data Fig. 8). Similarly, among the potential TISs in Arabidopsis seedlings identified by RiboTISH, their FDRs and predicted scores from our model were positively correlated (bottom panels in Fig. 7a). When we focused on RiboTISH-reported TISs in the monocots maize and rice, we also observed this correlation in the 5′ UTR-nonAUG TIS groups (left panels in Fig. 7b). Note that the sample sizes of the RiboTISH-reported 5′ UTR-AUG and CDS-AUG groups were small in some species, which might influence the results (Fig. 7a,b; Extended Data Fig. 8). These results demonstrate the feasibility and flexibility of our analytical pipelines and the utility of the TIS prediction model for identifying TISs from different plant tissues, as well as within and across species.

**Figure 7.**
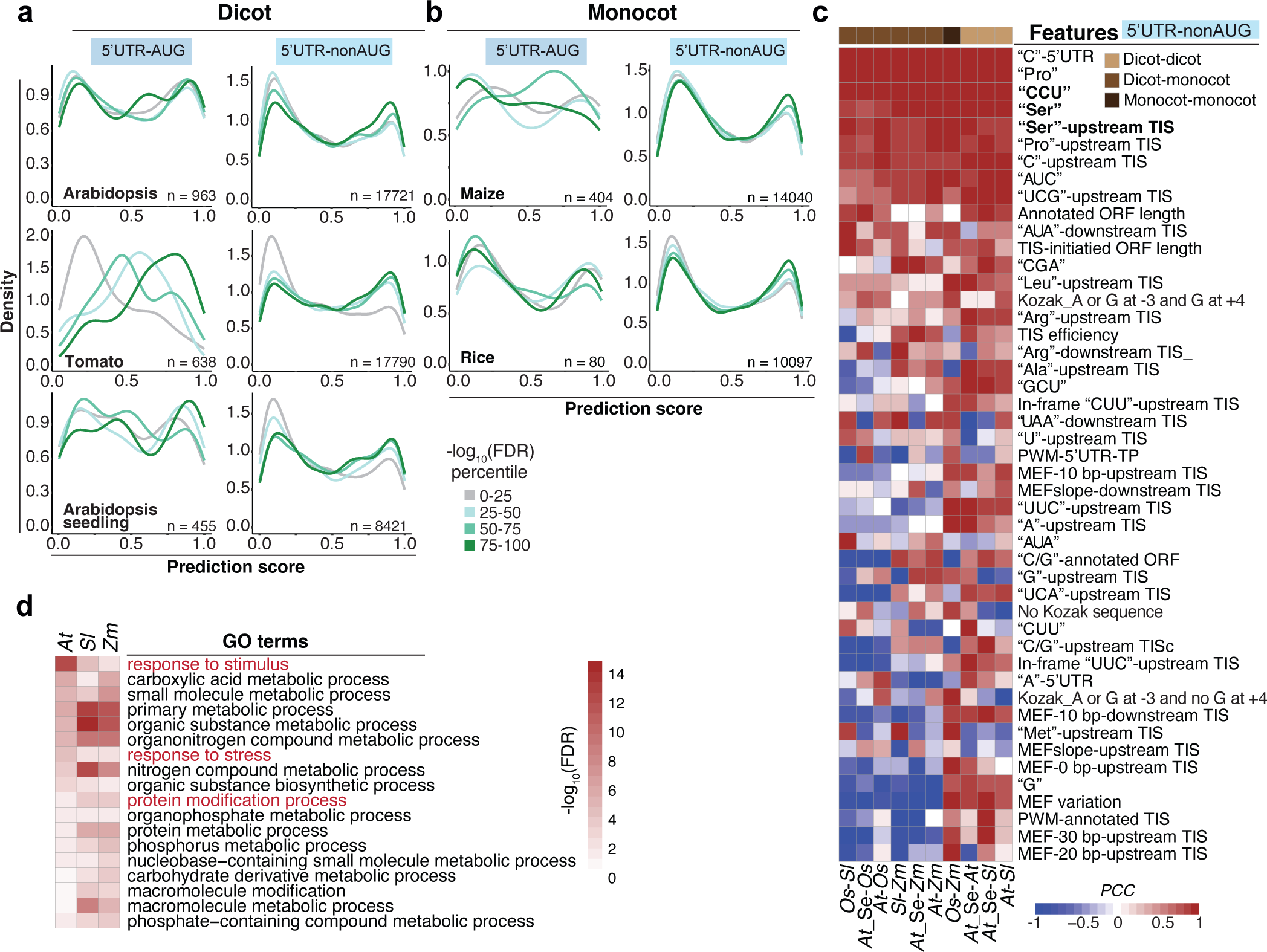
Prediction of monocot and dicot TISs and their characteristics. (a) Distribution of the TIS prediction scores generated by the Arabidopsis best models for the 5′ UTR-AUG and 5′ UTR-nonAUG TIS groups identified by the RiboTISH algorism and with RiboTISH-reported FDRs (FDR percentile in which 0-25 category includes TISs with lowest FDR values) using ribosome profiling datasets generated from dicot plants including Arabidopsis (suspension cells), tomato (leaves) and Arabidopsis seeding. (b) As indicated in (a), but for the datasets generated from monocot plants including maize and rice. (c) Pairwise Pearson’s correlation coefficient of the features among dicots (Arabidopsis (At), tomato (Sl), Arabidopsis seedling (At-Se) and monocots (maize (Zm) and rice (Os)) plants for the 5′ UTR-nonAUG TISs identified by the RiboTISH algorism. (d) GO biological process terms significantly enriched (FDR<0.05) in genes that have RiboTISH-reported TISs with top 10% highest prediction scores and with top 25% highest CG contents in the upstream 100-bp regions.

We calculated pairwise Pearson’s correlation coefficient (PCC) values to identify conserved features among different monocot and dicot plants. The contextual features related to “C” content, “Ser”, or “CCU” frequencies showed strong positive correlations (PCC > 0.8), suggesting that CU tracts are evolutionarily conserved features for TIS prediction (bold in Fig. 7c). The functions of genes with alternative TISs with the top 10% highest prediction scores and with frequent CU tracts (top 25% in highest-CU contents in its upstream 100-bp region) were associated with abiotic/biotic stress responses in all species examined, except for genes from the rice dataset, where no GO terms were enriched (Fig. 7d). Given the known role of CU-rich tracts in plant virus TIS activity (Fig. 5c)^35,49^, their validated function in promoting plant translation initiation (Figs. 5,7), and their association with stress-related plant genes (Fig. 7), CU tracts are likely important for triggering plant immunity and viral pathogenesis in both plants and plant viruses.

## Discussion

Gene annotation is critical for inferring gene family structure, evolution, and function and for decoding genomes from sequences to phenotypes, as it provides physical and biological contexts to an assembled genome sequence ^16^. The prevalence of unannotated protein-coding regions and the lack of a workflow for revising protein-coding gene annotations of existing plant genome references have limited fundamental discovery. Here, we addressed these issues by focusing on Arabidopsis and the agriculturally important tomato crop. Exploiting an ML pipeline, we discovered known Kozak motifs, novel CU-rich sequences, and the codons of TISs themselves with joint and distinct influences on alternative AUG and non-AUG TIS recognition (Figs. 1–4). We identified CU-rich elements in plant mRNAs that promote translation initiation (Fig. 5). We showed that evolutionarily conversed *cis*-regulatory signatures, particularly CU-rich sequences near TISs, were critical for accurate TIS prediction within and across plant species, as well as in humans and viruses (Figs. 1,2,3,7). Lastly, our models revealed hidden TISs based on mRNA sequences across monocots and dicots, thereby improving gene annotation in plants (Figs. 6,7). The translation of small or non-AUG TIS-initiated ORFs can expand proteome diversity and produce proteins with varied functions (Figs. 5,6 and Extended Data Fig. 6)^6,9,10,12^. Alternatively, some non-canonical ORFs act as *cis*-regulatory units to interfere with the translation of the main ORF in a transcript^50,51^. Thus, the coding regions or peptide compositions of these non-canonical ORFs may not be critical or conserved across plants. Additionally, since unannotated TIS-initiated ORFs tend to be short, the use of computational approaches is not very effective due to low conservation scores, highlighting the limitation of relying on sequence conservation in protein-coding regions for ORF annotation^52^. Our pipeline generates global estimates of the TISs used *in vivo*, providing protein-coding information for genomic sequences and facilitating efforts to bridge genotype to phenotype in plants.

Different ML pipelines have been employed to predict TISs in human and mouse cells^30,53,54^. These pipelines robustly identified TISs in mammals and revealed significant sequence features that provided information for predicting TISs^30,53^. In addition to known biological features such as the Kozak sequence and the sequences nearby flanking TISs, several important contextual features also contribute to TIS recognition mechanisms. For example, the number of upstream AUG/Ser residues contributed to the models with high importance scores in both humans and mouse (Fig. 2l)^30,53^. CU-rich tracts regulate TIS activities in humans and plant and human viruses (Figs. 3k,l and 5c)^35–37^. Consistent with *in silico* predictions, plant protein abundance decreased when the contextual features of CU-rich tracts were mutated (Fig. 5). Thus, given the similarity of the contextual features required for TIS prediction/activities in humans, mouse, viruses, and plants (Figs. 2,3,5), it is likely that these features are biologically meaningful for the regulation of TIS selection in eukaryotes rather than being merely the result of sequencing or experimental bias. The codon bias of TIS sites was also an important feature for plant TIS prediction (Extended Data Fig. 4), which is in line with findings for mammalian TIS prediction^53^ and the observation that AAG and AGG are generally poor start codons, as revealed using *in vitro*/*in vivo* reporter assay systems^4^. These observations demonstrate that both contextual features and TIS codon preference play important roles in TIS recognition in eukaryotes.

The polypyrimidine CU-tract element enhances translation initiation of preferred start sites via cap-independent and IRES-mediated translation in plant viruses. These CU-tract regions provide complementary interactions between the CU-rich regions of viral mRNAs and the conserved regions of 18S rRNAs within 40S small ribosomes, as reported for Tobacco etch virus, Blackcurrant reversion virus, Turnip Crinkle Virus, and Triticum Mosaic Virus^35,39,49,55^. A systemic high-throughput screen of IRES elements in humans and human viruses revealed short motif sequences including the known viral “UUCCUUU” and “UACUCC” IRES elements and novel short C/U-related sequences (such as “CCCUCUU” and “UUCCUU) that can base-pair with 18S rRNAs within a scanning ribosome to enhance cap-independent translation^36^. However, much less evidence is available for their regulatory roles in plant mRNA translation, with only a single report showing that a 100-bp CU-rich region within the 5′ UTR of *OsMac1* in rice promotes translation initiation^56^. Base-paring interaction between plant mRNAs and plant 18S RNAs is critical for translation initiation efficiency, as observed in the short 5′ UTRs of Arabidopsis ribosomal protein S18C and a plant translation enhancer element with the active plant ribosomal 18S RNA complementary sequences^41,57^. In line with these findings for individual genes, our studies systemically and experimentally characterized the broad influence of CU-rich sequences in plant TIS recognition across different plant species, likely via interaction with plant 18S RNAs, and highlighted a conserved translational strategy that modulate initiation site recognition in eukaryotes (Figs. 3,5,7). In addition, since plant viruses hijack the host translational machinery and use non-canonical translational strategies for their protein production^58^, it is likely that both plants and plant viruses, which have evolved to drive host factors, exploit similar efficient sequence elements to facilitate the translation of individual proteins. Moreover, genes with alternative TISs were associated with abiotic/biotic stress responses (Fig. 7e). It will be interesting to investigate how plants and plant viruses use these CU-rich tracts for protein synthesis and which translated proteins result in plant immunity.

Protein synthesis is an essential but costly program, accounting for 30%–50% of energy consumption in cells^59,60^. The conservation of TIS recognition mechanisms reflects their critical and universal roles in controlling mRNA translation, which is indispensable for all organisms. Whereas translation initiation can occur at non-AUG sites, initiation at these sites is tightly controlled via a delicate recognition mechanism (Figs. 1–4) that minimizes noisy translation initiation. Given that protein production is tightly regulated *de novo* by *cis*-regulatory elements near TISs, our findings of sequence context features point to the utility for precisely controlling protein production circuits in plants. Indeed, upstream ORF-mediated translation has been used to engineer disease resistance in Arabidopsis and rice^61^.

Kozak sequences and the sequences nearby flanking TISs are well-known *cis*-regulatory signatures that enhance initiation at either AUG or non-AUG start codons^19,32^. Here, we showed that Kozak motifs are preferentially associated with 5′ UTR-nonAUG and CDS-AUG, but not 5′ UTR-AUG TISs (Figs. 2,3; Extended Data Fig. 3). How can ribosomes recognize these *cis*-regulatory sequences when scanning mRNAs? Multiple *trans*-acting factors including (but not limited to) eukaryotic translation initiation factors (eIFs) play vital roles in selectively regulating TIS efficiency^4^. Genetic analyses of yeast (*Saccharomyces cerevisiae*) eIFs have revealed the distinct regulatory roles of eIF1, eIF1A, and eIF2 in recognizing AUG and non-AUG TISs^4^. There is also structural evidence that interactions among different eIFs and the initiator transfer RNA (Met-tRNAi) and mRNA templates affect base-paring between anticodons and codons or changes in the conformation of the preinitiation complex at AUG and non-AUG sites^18^. In addition, the abundance of eIFs is often regulated by stress conditions, and these proteins in turn initiate alternative translation events and subsequently affect the production of proteins that regulate cell survival under stress conditions^4^. Together, these findings suggest that *cis*- and *trans*-regulatory factors orchestrate a regulatory network for TIS selection and allow sophisticated co-option for condition-dependent translational regulation.

Our TIS prediction models identified informative *cis*-regulatory features in Arabidopsis and tomato, revealing the mechanistic basis of alternative TIS recognition across dicot and monocot plants. The evolutionary similarities of plant TIS recognition principles highlight the feasibility of applying TIS prediction models to crop species with little experimentally derived gene information. Integrating these prediction models into existing bioinformatics tools would leverage the power of protein-coding gene annotation pipelines across diverse plant species. Moreover, these *cis*-regulatory features will facilitate efforts to reveal the associations between genetic variation, gene expression, and phenotypic diversity, ultimately enhancing the discovery of functional genes, especially in information-poor crop species.

## Materials and Methods

### Generation of the true-positive (TP) and true-negative (TN) translation initiation sites (TISs)

The TP TISs were defined as TISs with significant translation initiation activities and were identified by analysis of the LTM- and CHX-treated ribosome profiling datasets using a TIS-finding pipeline as described previously^62^. The LTM- and CHX-treated ribosome profiling datasets generated from Arabidopsis suspension cells and tomato leaves were retrieved from Gene Expression Omnibus database (GSE88790 and GSE143311)^6,10^ and used to profile the translating ribosome positions on transcripts as performed previously^6^. The TIS-finding pipeline^62^ with default parameters was used to call the TIS peaks by employing the LTM and CHX datasets to identify the TISs used *in vivo*, referred as the TP TISs. Briefly, a zero-truncated negative binomial distribution (ZTNB) was performed to statistically model (1) the background distribution of LTM + CHX pooled counts in genomes (i.e., all non-zero positions with LTM + CHX pooled counts across all transcripts) to obtain a global threshold with a *p*-value < 0.05 and (2) the background distribution of LTM + CHX pooled counts and CHX counts in transcripts with more than 50 positions with non-zero counts to obtain the local *p*-values for each position on a transcript. The candidate start site examined was called a TIS peak based on the following criteria: (1) an LTM count and CHX count both >0; (2) an LTM + CHX pooled count > the global threshold; (3) a local *p*-value of the LTM + CHX pooled counts <0.01 and 1000-fold higher than the local *p*-value of the CHX counts at the same location or a local *p*-value of the LTM + CHX pooled counts less than 10^−7^.

To generate reliable TN TIS datasets, we focused on AUG and near-cognate codons and only searched for the TN sites in transcripts with identified TP TISs. The candidate codon sites were referred as to TN sites based on the following criteria: (1) LTM and CHX counts both = 0; (2) the site is located in the upstream region of the most downstream TP TIS on the same transcript as descripted previously^30^.

### Identification of putative cis-elements for predicting TISs

To search for the putative *cis*-regulatory elements associated with Arabidopsis AUG/non-AUG TIS activities, an enrichment-based *k*-mer (oligomer with the length of *k*)-finding pipeline was used as described previously^26^. Briefly, all possible *k*-mer sequences (*k* ≥ 4) were examined for significant enrichment in the 200-bp transcript regions centered on a given TIS between the TP and TN TISs. For the shorter *k*-mer sequences that were perfectly matched to the longer *k*-mer sequences, only the one with higher enrichment significance (i.e., the lower *p*-value) was referred to as a putative *cis*-regulatory element in the downstream TIS prediction analyses.

### Feature collection

Feature collection was based on a previous report^30^ with slight modifications unless specified otherwise. The definition, generation and slight modification of known/open-reading frame (ORF)/contextual/TIS codon usage feature categories are described below.

### -- Known features

(1) Position-weight-matrix (PWM): multiple PWM-related features representing the relationship between the flanking sequence context and the TIS translational efficiency were determined as follows. First, the features “PWM_TP_” and “PWM_annotated_” were determined by a PWM matrix generated based on the flanking sequences (positions −15 to +10) of a given TP TIS group and of the annotated TISs with *in vivo* translation initiation activity. Second, the feature “Noderer translational efficiency”, representing the relationship between the flanking sequences (position - 6 to +5) and the TIS translational efficiency in mammalian cells, was derived from a previous report^20^. In brief, all possible flanking sequence (positions −6 to +5) contexts around the AUG translational start were identified as features.
(2) Kozak sequence context: The Kozak sequence context was discretized into strong (A or G at - 3 and G at +4), intermediate (A or G at −3 and no G at +4), weak (no A and no G at −3 and G at +4), and no Kozak context. These categories were presented as the values 1 (no), 2 (weak), 3 (intermediate), and 4 (strong).

### -- ORF features

(1) ORF length: arbitrary start sites in the mRNA sequence, the lengths of the TP/TN TIS- and annotated TIS-initiated ORFs, and the A/T/C/G mononucleotide contents in their upstream regions were considered.
(2) Minimum free energy (MFE) of mRNA secondary structure: the MFEs in the 80-bp regions centered on TISs were calculated using the RNAfold program^63^ in a sliding window with a 20-nt window size and a 10-nt step size. In addition, to summarize the magnitudes of the MFE difference, the 20-bp upstream and downstream regions flanking the TISs were also computed by normalizing the MEF values of the region at positions −20 to +0 to those of the regions at positions −10 to +10 and the MEF values of the region at positions +10 to +30 to those of the regions at positions +0 to +20.

### -- Contextual features

We counted the frequency of all possible *k*-mers of length *k* = 1 (position-specific *k*-mers) and *k* = 3 (codon and respective amino acid *k*-mers) in a window from −99 to +99 around the start site. The *k*-mers were defined as all possible combinations of subsequence of length *k*, given one of the four nucleotides A, U, C, and G. The in-frame and out-of-frame *k*-mers as well as *k*-mers upstream and/or downstream of the start site were considered. In addition, we also considered the frequency of all possible amino acids with length =1 and 2 in the 99-nt regions downstream of the start site and within a TIS-initiated ORF. These generated 2626 contextual features in total.

### -- TIS codon usage

The TIS codon usage of all 64 codons was determined as described previously^53^. Briefly, for a given codon, the proportion of the target codon sites among all the identified TP TISs were normalized to the proportion of the target codon sites among all codon sites found in the transcript regions of all annotated genes; then the corresponding log2 ratio was computed and referred to as the feature value of the TIS codon usage bias.

### Feature selection and data imputation

The size of the TN/TP dataset was balanced to contain the same number of TN and TP sites by randomly under-sampling from the larger dataset. We applied Wilcoxon-rank sum test and Bonferroni correction for all features to test for the statistical significance of differences between TP and TN sites. We then calculated Pearson correlation coefficients among the contextual and ORF features. The 50 most significant (with smallest adjusted *p*-value) and uncorrelated (r < 0.7) contextual features and the uncorrelated (r < 0.7; adjusted *p*-value < 0.01) ORF features were kept for the model training step. All features were normalized and scaled to ensure comparability. We repeated this selection 10 times to evaluate the model robustness for each TIS group.

### Construction and evaluation of predictive machine learning models

An ML pipeline described previously^24,64^ was retrieved from GitHub (https://github.com/ShiuLab/ML-Pipeline). Briefly, we used scikit-learn (v0.24.2) in Python (v3.7.0) to train and test the models. For each TIS group, we split balanced data into training (70%) and testing (30%) sets and tested four classification methods: Random Forest (RF), Support Vector Machine (SVM), Logistic Regression (LR), and Gradient Boost (GB). We used ten-fold internal cross-validation to select the optimized hyperparameters. F1 score was used to select the best model for each TIS group. Note that to have TIS-predictive features representative in a given species, only the features used in the best model with frequency >=7 in the 10 randomly balanced datasets were included in Fig. 1g. To comprehensively reveal the features contributing to the best model of predicting TISs, the features used at least in one of the 10 randomly balanced datasets were included in Fig. 2a,c,e,g and Extended Data Fig. 2c,e,g,i.

### Generation of the candidate TIS-initiated protein expression constructs with mutations and expression tags

The best models generated (Fig. 1e and Extended Data Fig. 1e) were used to compute the possibility of a given triplet being classified as a TP. The triplets with the prediction scores higher than the threshold of classifying TP/TNs used in best models (Fig. 6a and Extended Data Fig. 7) were selected for further functional validation.

The introduction of protein expression constructs into tobacco using an Agrobacterium-mediated transient expression system and detection of expression were performed as described previously^6^ with some minor modifications. Briefly, the 5′ UTR and CDS fragments of the gene with a given target TIS site ranging from the upstream 100 bp to the downstream 9 nt (including the target TIS site) were amplified by PCR and fused with a reporter gene encoding GFP, 10xMYC, or 3xFlag, which were used previously^6,65^. All mutagenesis of the tested sites of genes was performed using synthetic primers listed in Supplementary Table 3.

### Quantitative reverse-transcription PCR analyses

Total RNAs (2–3 μg) were extracted from tobacco leaves using PureLink Plant RNA Reagent (Invitrogen, #12322012) and used for cDNA synthesis with M-MLV Reverse Transcriptase (Invitrogen, #28025013). The transcript quantification of the target genes containing TISs of interest and *Ubiquitin 3*^66^ (as an internal control) were detected via qPCRBIO SyGreen Mix (PCR Biosystems Ltd.) and analyzed on a BioRad Real-Time PCR System. Primers used are listed in Supplementary Table 3.

### Detecting potential TISs using RiboTISH software and generating their prediction scores

The public CHX-treated ribosome-profiling datasets in Arabidopsis (suspension cells), tomato, maize and rice were retrieved from Gene Expression Omnibus database (GSE88790, GSE143311)^6,10^ and NCBI Sequence Read Archive (SRA) database (PRJNA523300 and SRP052520)^46,47^. To assess the model performance between different plant tissues in Arabidopsis, the CHX-treated ribosome profiling datasets of 6-day-old Arabidopsis seedling were generated as described previously^6^. The raw reads were trimmed and mapped as described in previous respective studies^6,46,47^ and then input into RiboTISH algorism^48^ with default setting and the additional parameters of “-longest -alt” for the identification of the TISs with FDRs (i.e., the BH correction q-value of frame test). The prediction scores for each RiboTISH-reported TIS were calculated in scikit-learn using the best model from Arabidopsis (suspension cells). The genome version used in this study was Zm-B73-REFERENCE-NAM-5.0 and IRGSP-1.0 for maize and rice, respectively.

### Gene Ontology (GO) analysis

GO term enrichment analysis was performed with the Panther database^12^ using Fisher’s exact test to calculate the degree of enrichment with FDR for a multiple testing adjustment. Significantly enriched GO terms (FDR < 0.05) were visualized as heatmaps.

## Acknowledgments

We thank Mr. Te-Chang Hsu, Dr. Yao-Cheng Lin, and the AS-BCST Bioinformatics Core for high-performance computing services and Dr. Shin-Han Shiu and Dr. Ho-Ming Chen for critical reading and Dr. Melissa Lehti-Shiu for English editing of this article.

## Author contributions

T.-Y. W. designed the research, performed bioinformatics analyses, and wrote the paper. K.-J. C. performed bioinformatics analyses. Y.-R. L. performed the experimental analyses. D. U. revised the paper. M.-J. L. conceived/designed the research, performed bioinformatics analyses, and wrote the paper.

## SUPPLEMENTARY MATERIALS

***Extended Data Figure 1. The characteristics of the identified translation initiation sites (TISs) in Arabidopsis and tomato.***

(a) The positional distributions of the identified TISs mapping to different genic regions of transcripts including the 5’ UTRs, annotated AUG TIS sites, CDS and 3’ UTRs. Background: the locations of all 64 codons in transcripts. (b) The codon compositions of the identified TISs located in the 5′ UTRs and CDS. Near-cognate: the codons that differ from AUG by one base. Others: codons other than AUG and near-cognate codons. (c) The codon compositions of the identified TISs containing AUG and near-cognate codons for the Arabidopsis TISs located in the 5′ UTRs (left) and CDS (right). Background: the codon compositions based on all transcript sequences. (d) As described in (c), but for the identified TISs in tomato. (e) The prediction performance (represented as F1 scores) when all features (i.e., the Known/ORF/contextual ones) were applied in predicting the four tomato TIS groups. Circle: the performance of a model with a given combination of model parameters in a randomly balanced TP and TN dataset. Black line: mean of the F1 scores for a given ML algorithm. Arrow: the best model (i.e., for the ML algorithm with highest mean performance, the model with highest F1 score). Dashed line: the baseline performance expected by random guessing.

***Extended Data Figure 2. The features that were most informative for predicting TISs in Arabidopsis and tomato.***

(a,b) Feature Elimination analyses of selecting the top 10 features with the highest importance to the performance of the best model. Boxplots present the F1 scores generated from 10 balanced datasets (black dots) randomly chosen from the four TIS groups in Arabidopsis (a) and tomato (b); the median values, the first and third quartiles, and whiskers of maximum and minimum values are shown. (c) Comparison of the importance scores derived from the best model and the statistical significance of differences (-log_10_(FDR), determined by Wilcoxon signed-rank test) between Arabidopsis CDS-AUG TPs and TNs for the features used in the best model. Rho: Spearman’s rank correlation coefficient. The black line indicates the fitted linear regression line, and the gray area indicates the 95% confidence level interval. (d) The means of the feature values in the Arabidopsis CDS-AUG TP and TN datasets (left) and the frequency of the Feature Elimination-determined top10 features (ranked using their importance revealed in (a)) identified in 10 randomly balanced datasets (left). The rank and frequency indicate the importance of a given feature in the prediction model and their robustness using 10 randomly balanced datasets. The features with frequency >7 within the top 10 are shown. Orange indicates the TIS group with the higher feature value. (e-j) As described in (c,d), but for the Arabidopsis CDS-nonAUG TIS group (e,f), the tomato CDS-AUG TIS group (g,h) and tomato CDS-nonAUG TIS group (i,j).

***Extended Data Figure 3. The C/U nucleotide compositions and the flanking sequences of CDS TISs.***

(a,b) Sequence logo plots showing the differential enrichment of A/U/C/G nucleotides between TPs and TNs in the region 15-bp upstream and 13-bp downstream of the CDS-AUG and CDS-nonAUG TISs, represented as the log2 ratio of the site frequencies between TPs and TNs, in Arabidopsis (a) and tomato (b). (c,d) Position-weight matrix (PWM) scores showing the sequence similarity of the TIS-flanking regions between TPs (brown)/TNs (gray) and annotated TISs for the Arabidopsis (c) and tomato CDS TISs (d). PWM_annotated TISs_ matrix was determined based on the annotated TISs with *in vivo* translation initiation activities (See Methods). P-values derived from Mann–Whitney U test were used to evaluate the significance of differences of PWM scores between the TP and TN datasets.

***Extended Data Figure 4. The inclusion of TIS codon usage bias as a feature improves prediction of non-AUG TISs in Arabidopsis and tomato.***

(a) Comparison of the prediction performances with (red) and without (gray) TIS codon usage bias as a feature for the 5′ UTR-nonAUG and CDS-nonAUG TIS groups in Arabidopsis and tomato. As described in Fig. 1E, but for the median F1 scores of the ML algorithms with the highest performance. (b) Box plots showing how the TIS codon preference feature affects the proportion of the mis-predicted TISs in the TNs (TN→TP; light brown, left) and TPs (TP→TN; dark brown, right) of the 5′ UTR-nonAUG TIS groups. (c) As described in (b), but for individual near-cognate codons. Arrows: the top two codons with the highest degree of difference in mis-prediction with/without using the feature of TIS codon usage bias (orange vs. green).

***Extended Data Figure 5. The C/U nucleotide compositions and the flanking sequences of Arabidopsis and tomato TISs.***

(a,b) Enrichment of sites with the indicated 3-mer sequences in the CDS-AUG and CDS-nonAUG TIS groups, represented as the log2 ratio of the site frequencies between TPs and TNs in the 180-bp region centered on TISs with a 10-bp window, in Arabidopsis (a) and tomato (b). (c-f,h) As described in (a,b), but for the “AUG” and “UUC” sequences (c,d), the A, U, C, and G mononucleotides (E,F) and the “UCUUC” and “UCUCU” sequences (h). (g) As described in Extended Data Fig. 2c, but for the putative RNA *cis*-elements in predicting the Arabidopsis CDS-AUG and CDS-nonAUG TIS groups. (i,j) Relationships between the number of C/Us (y-axis) and the enrichment of sequence occurrence (x-axis) for all the 5-mers in the Arabidopsis (i) and tomato (j) CDS-AUG and CDS-nonAUG TIS groups. The enrichment is represented as the log2 ratio of the median of the sequence occurrence between TPs and TNs in the 200-bp regions centered on TIS sites. Rho: Spearman’s rank correlation coefficient. Arrows: the putative *cis*-elements discussed in Fig. 4.

***Extended Data Figure 6. The in vivo initiation activities of mis-classified TISs revealed via ribosome profiling***

(a) Plots showing the LTM read counts of the indicated genic regions in two biological replicates for Solyc03g096920.3.1, which has a 5′ UTR-AUG TIS (uTIS, AUG; orange arrow) validated previously^6^. The gene model (bottom) with the UTRs (light gray boxes), annotated CDSs (dark gray boxes), introns (thin lines), annotated TIS (aTIS, black arrow), and the upstream 100-nt wild-type (WT) sequence or sequence with mutations of CT tracts (red) are shown. The upstream 100-nt region was divided into 6 subregions whereas the leftmost one is the 1^st^ subregion as indicated in Fig. 5d. (b) As described in (a), but for Solyc07g052600.3.1, which has a 5’UTR-AUG uTIS site (uTIS, GUG; black arrow) validated previously^6^. (c) As described in (a), but for Solyc06g76770.3.1, which has a mis-classified 5′ UTR -AUG site (uTIS, AUG; orange arrow). There were statistically significant TIS signals for this TIS in replicate #1 but not in replicate #2 because of the low read counts in replicate #2. Although the reads did not pass the detection threshold, the prediction score of 0.94 passed the prediction threshold of 0.41. (d) As described in (a), but for Solyc06g009750.3.1, which has a mis-classified 5’ UTR-CUG site (uTIS, CUG; orange arrow). The signals for this TIS were statistically significant in replicate #2 but not in replicate #1, because of a low read count in replicate #1, which did not reach the detection threshold, but the prediction score of 0.81 passed the prediction threshold of 0.44. (e) As described in (a), but for Solyc04g76110.3.1 with the indicated mis-classified 5′ UTR-AUG site (uTIS, AUG; orange arrow). The TIS signals were significant in replicate #2 but not in replicate #1, probably because of the low precision of RPF read mapping at the uTIS, but the prediction score of 0.99 passed the prediction threshold of 0.41. (f) As described in (a), but for Solyc05g050510.3.1, which has the indicated mis-classified 5′ UTR-nonAUG TIS (uTIS, GUG; orange arrow). There were no significant LTM signals for this TIS in both replicates, but the prediction score of 0.61 passed the prediction threshold of 0.44. (g) As described in (a), but for Solyc11g039830.3.1, which has the indicated mis-classified CDS-AUG TIS site (dTIS, in-frame AUG; orange arrow). The signals for this TIS were statistically significant in replicate #2 but not in replicate #1 because of the low read count in replicate #1, which did not pass the detection threshold, but the prediction score of 0.66 passed the prediction threshold of 0.49. The annotated AUG TIS-encoded protein isoform, but not the AUG dTIS-encoded one, has a predicted mitochondria-targeting signal (red).

***Extended Data Figure 7. The feature characteristics of the tomato mis-classified TN TISs***

(a,b) Prediction score distribution of the CDS-AUG and CDS-nonAUG TIS groups in Arabidopsis (a) and in tomato (b). The mean threshold (dashed lines) for classifying the TN→TN (the TN TISs predicted as TNs, gray) and TN→TP (the TN TISs predicted as TPs, light brown) groups derived from the models on the10 randomly balanced datasets (as indicated in Fig. 1) are shown. (c,d) Dot plots show the frequency (y-axis) of a given feature used for TIS prediction in 10 randomly balanced datasets and the feature enrichment (FDR, x-axis) between the TN→TN and TN→TP groups for the TIS groups indicated in (a,b). The red line represents the threshold (frequency ≥ 7) of important features as indicated in Fig. 2. (e,f) Violin plots show the feature value distributions for the features that were most enriched in (c,d) for the TN→TN (gray) and TN→TP (light brown) groups, indicated in (a,b), and the TP→TP (the TP TISs predicted as TPs, dark brown) group. The red dot represents the median value.

***Extended Data Figure 8. Prediction of monocot and dicot TISs using transfer learning***

Distribution of the TIS prediction scores generated by the Arabidopsis best models for the CDS-AUG and CDS-nonAUG TIS groups identified by the RiboTISH algorism and with RiboTISH-reported FDRs (FDR percentile in which 0-25 category includes TISs with lowest FDR values) using ribosome profiling datasets generated from dicot plants including Arabidopsis (suspension cells), tomato (leaves) and Arabidopsis seedling and monocot plants including maize and rice.

***Supplementary Table 1. The mean F1 scores showing the performances of models predicting translation initiation sites (TISs) based on the indicated features.***

***Supplementary Table 2. The mean feature value (with standard deviation) of true and false start sites, the significant differences of feature values between true and false start sites, linear regression coefficients for the features that were used in the best human linear regression model (Reuter et. al., 2016).***

***Supplementary Table 3. List of primers used in this study.***

